# High-resolution Expression Profiling of Selected Gene Sets during Plant Immune Activation

**DOI:** 10.1101/775973

**Authors:** Pingtao Ding, Bruno Pok Man Ngou, Oliver J. Furzer, Toshiyuki Sakai, Ram Krishna Shrestha, Dan MacLean, Jonathan D. G. Jones

## Abstract

Sequence capture followed by next-generation sequencing has broad applications in cost-effective exploration of biological processes at high resolution [1, 2]. Genome-wide RNA sequencing (RNA-seq) over a time course can reveal the dynamics of differential gene expression. However, in many cases, only a limited set of genes are of interest, and are repeatedly used as markers for certain biological processes. Sequence capture can help generate high-resolution quantitative datasets to assess changes in abundance of selected genes. We previously used sequence capture to accelerate *Resistance* gene cloning [1, 3, 4], investigate immune receptor gene diversity [5] and investigate pathogen diversity and evolution [6, 7].

The plant immune system involves detection of pathogens via both cell-surface and intracellular receptors. Both receptor classes can induce transcriptional reprogramming that elevates disease resistance [8]. To assess differential gene expression during plant immunity, we developed and deployed quantitative sequence capture (CAP-I). We designed and synthesized biotinylated single-strand RNA bait libraries targeted to a subset of defense genes, and generated sequence capture data from 99 RNA-seq libraries. We built a data processing pipeline to quantify the RNA-CAP-I-seq data, and visualize differential gene expression. Sequence capture in combination with quantitative RNA-seq enabled cost-effective assessment of the expression profile of a specified subset of genes. Quantitative sequence capture is not limited to RNA-seq or any specific organism and can potentially be incorporated into automated platforms for high-throughput sequencing.

## RESULTS AND DISCUSSION

In previous work, we investigated changes in *Arabidopsis thaliana* defense gene expression in response to a bacterial effector after recognition via nucleotide-binding leucine-rich-repeat intracellular immune receptors (NLRs). Specifically, we delivered the *Ralstonia solanacearum* effector PopP2, and studied responses to its recognition by the RPS4/RRS1-R intracellular immune receptor complex [9]. We defined a subset of early response genes (ERGs) particularly responsive to NLR activation (Fig S1A, Table S1 and S2). Expression of ERGs can be induced by both cell-surface receptors and NLRs, but more rapidly and strongly induced when both classes of receptors are activated (Fig S1A). NLR-dependent ERG upregulation was first observed at four hours post-infiltration (4 hpi) (Fig S1B, C). To assess the roles of immune components during ERG activation, we measured ERG transcripts in selected immune-deficient mutants compared to wild type (wt). Since these studies involved multiple replicates, mutant backgrounds and treatments, we applied complexity reduction via sequence capture to reduce sequencing costs.

We selected investigated 35 ERGs, and also 17 non-ERGs as controls, based on their transcriptional regulation patterns (Fig S1A, Table S2) [9]. The ERGs include genes that are important for conferring full resistance to various plant pathogens, and are involved in the biosynthesis of phytohormones, salicylic acid (SA) and pipecolic acid (Pip), including *ICS1*, *EDS5, PBS3, FMO1* and genes that encode the transcription factors (TFs) WRKY51 and SARD1 [10–16, 17]. Non-ERG control genes include *UBQ10* and *ACT7*, as well as late immune response genes [9], such as *PR1*, which is known to be activated by elevated SA [18]. We included full-length gene loci as templates for the capture bait design, spanning gene bodies (introns included) and putative promoters and terminators (Fig 1A). For promoters and terminators, we either defined them based on the intragenic sequence region between the coding sequence (CDS) of the target gene and the CDS of the immediate neighboring genes (<4,500 base pairs, or bps), or used 4,500 bps upstream of the start codon or downstream of the stop codon as their promoters or terminators, respectively (Fig 1A). This was to minimize the loss of any important sequence information: some genes might need longer intragenic regions to be fully functional. All sequence templates were designed using the gene coding strand (Fig 1A).

**Figure 1.**
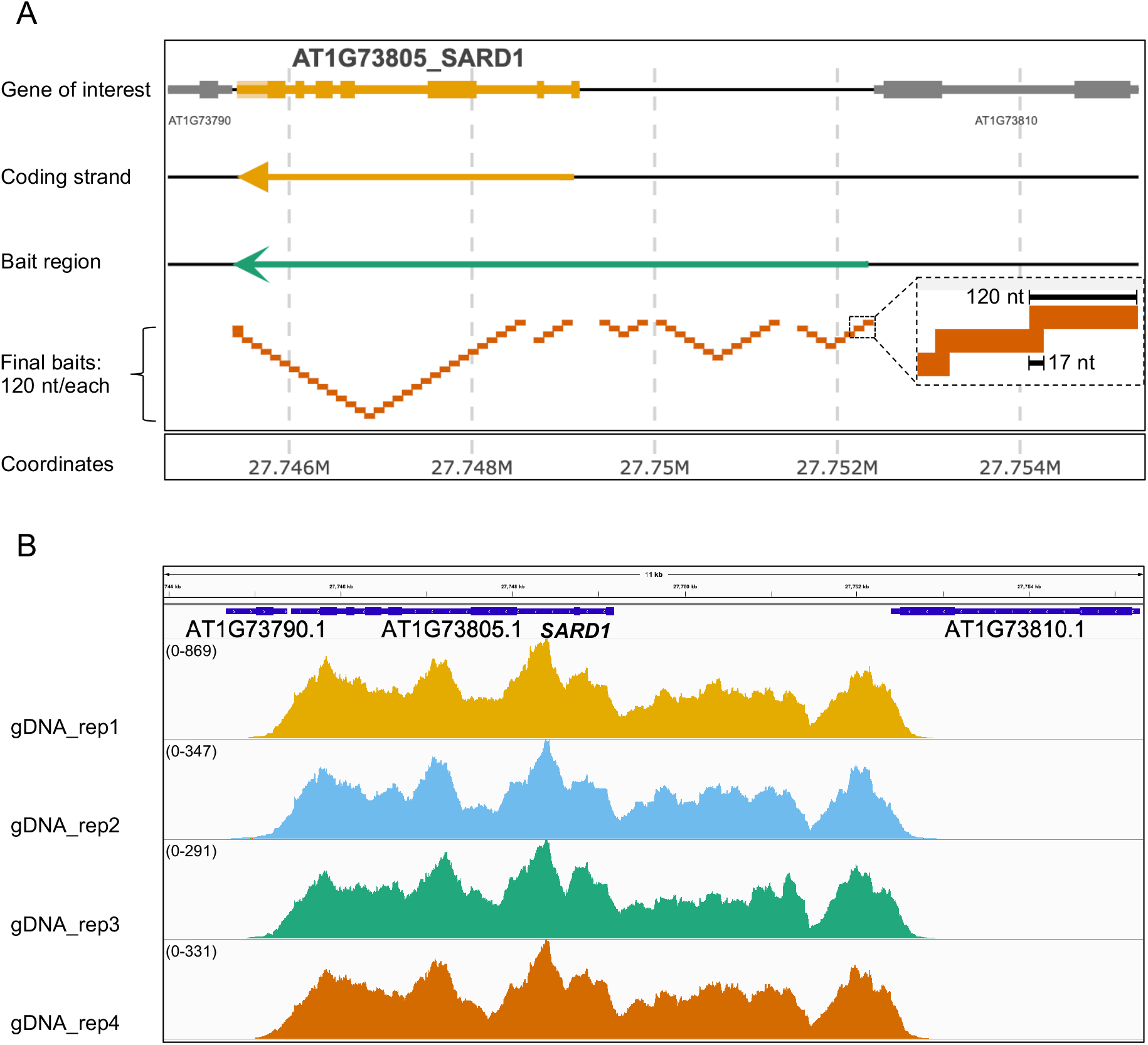
CAP-I Bait Design and Validation. (A) Visualization of bait design on one of CAP-I gene loci, *SARD1*. Using GFF file, here we present the genome organization of one CAP-I gene locus, *SARD1*. Top row shows the annotated exons and introns and intragenic regions of CAP-I gene locus and neighboring gene loci. Second row show the direction of the coding strand, here *SARD1* coding is on the reverse strand. The third row shows the orientation and the region that covers *SARD1* loci and putative promoter and terminator. The fourth strand shows the final non-redundant baits we designed and how they are mapped to the CAP-I target gene locus. The final baits are 120 nucleotides (nt) in length with 17 nt overlap for tilling. (B) Trial run of CAP-I-seq reads from genomic DNAs mapped to *SARD1* locus and visualized in a genome browser. Illumina sequencing reads of genomic DNA (gDNA) with four biological replicates in one CAP-I capture shows 100% coverage on all CAP-I gene loci including *SARD1*. See also Figure S1, Table S1 to S3.

After computationally extracting sequences from all 52 gene loci, we used our bait design pipeline to design a bait library (Fig 1A and Fig S2A). We synthesized a set of 20,000 120-mer single-strand RNA probes (Fig 1A), which contains 2219 unique probes with 17-nucleotide tiling and covering ~ 260 kb of the corresponding *Arabidopsis* genome regions (Fig S1A). We named this library as ‘Capture I’ (CAP-I) for studies of plant innate immunity. To test the efficiency of CAP-I for sequence capture, we performed one capture with libraries generated from *Arabidopsis* genomic DNA for NGS. We found all gene loci have 100% breadth of coverage (Fig 1B and Table S3), showing that CAP-I enables capture of targeted sequences (Fig 1B). The pipeline generated one set of redundant baits in the region between two adjacent genes (Fig S2B), which could be condensed to provide additional capture capacity.

We then tested if CAP-I can be used in RNA-seq to assess quantitative changes in ERG transcripts. We used *Arabidopsis thaliana* accession Col-0 as wt, and also investigated seven selected mutants in Col-0 (Fig S2C). Resistance to *Ralstonia solanacearum* 1 (RRS1)-S and RRS1B are NLRs of bacterial effector AvrRps4, and they function together with their paired NLRs Resistant to *Pseudomonas (P.) syringae* 4 (RPS4) and RPS4B, respectively [19]; a *rrs1-3 rrs1b-1* mutant loses AvrRps4 responsiveness. EDS1 (the included mutant is *eds1-2*) is required for immunity mediated by Toll/Interleukin-1 Receptor/Resistance (TIR)-NLRs like RRS1 and RPS4 [20]. *SID2* (the included mutant is *sid2-2*) encodes the enzyme ICS1, which is required for the biosynthesis of defense-related phytohormone, SA [10, 21]. SARD1 and its homolog Calmodulin-binding Protein 60-like g (CBP60g) are master TFs required for transcriptional regulation of genes involved in pathogen associated molecular pattern (PAMP)-triggered immunity (PTI), effector-triggered immunity (ETI) and systemic acquired resistance (SAR) [13, 14, 22–24]. MYC2 and its homologs MYC3 and MYC4 are basic helix-loop-helix TFs (the included mutant is *myc2 myc3 myc4*) required for jasmonic acid (JA)-mediated resistance against bacteria [25]. TOPLESS (TPL) and its homologs TPL-related 1 (TPR1) and TPR4 (the included mutant is *tpl tpr1 tpr4*) are putative transcriptional co-repressors required for full resistance against the bacterium *P. syringae* pv. *tomato* DC3000 (hereafter DC3000) and DC3000 expressing AvrRps4 but not DC3000 expressing AvrRpt2, an effector recognized by RPS2, a non-TIR-NLR [26]. Phytoalexin Deficient 4 (PAD4), Ethylene-insensitive protein 2 (EIN2), Delayed Dehiscence 2 (DDE2, encoding an allene oxide synthase involved in jasmonic acid synthesis) and SID2/ICS1 (the included mutant is *pad4-1 ein2-1 dde2-2 sid2-2*) are proteins that are involved in different but interacting sectors in immune signaling networks [27].

Previously, we have defined the response induced by the bacterium *P. fluorescens* (Pf0-1 EtHAn strain) carrying a mutant effector PopP2^C321A^ (Pf0-1:PopP2^C321A^) as ‘PTI’ mediated by cell-surface Pathogen Recognition Receptors (PRRs) [9]. The Pf0-1 strain carrying wt PopP2, recognized by RRS1-R/RPS4, triggers an additional ETI response that we designate ‘PTI + ETI’. Here, we used Pf0-1:AvrRps4 or Pf0-1:AvrRpt2 to induce ‘PTI + ETI’. The responses induced by Pf0-1:AvrRps4 or AvrRpt2 are named as ‘PTI plus TIR-NLR-mediated ETI’ (PTI + t-ETI) and ‘PTI plus CC-NLR-mediated ETI’ (PTI + c-ETI), respectively (Fig 3C). In addition, Pf0-1 carrying the mutant effector AvrRps4^KRVY135-138AAAA^ (Pf0-1:AvrRps4^KRVYmut^) was included as ‘PTI’. We also included leaves infiltrated with buffer only, as a mock treatment, and no treatment on wt plants as an untreated control (Fig S2C). ERGs began to show significant upregulation in their transcripts at 4 hpi of Pf0-1:PopP2 compared to Pf0-1:PopP2^C321A^ [9], so we collected our samples at 4 hpi for all treatments. For each combination of genotype and treatment, we collected 3 biological replicates; 99 samples in total (Fig S2C). We extracted RNAs from these samples and generated cDNA libraries. Each library was barcoded with custom index primers. In addition, we added genomic DNA libraries in the final multiplexed library as spike-in controls for sequence capture. We applied one reaction of CAP-I baits to capture the multiplexed libraries before sequencing.

After demultiplexing, we retrieved single-end reads for each individual library. We mapped the reads to CAP-I target gene loci and assessed the mapping efficiency. We observed 100% breadth of coverage of full-length transcripts for all gene loci except for *AT4G28410*, which encodes Root System Architecture 1 (RSA1). *RSA1* is specifically expressed in *Arabidopsis* root tissue, and all our samples are leaf tissues, so *RSA1* served as a good negative control for contamination introduced at any steps of library preparation and sequencing. Since no reads from 99 cDNA libraries of RNA-CAP-I-seq mapped to the *RSA1* locus while 100% breadth of coverage in *RSA1* locus occurred in the gDNA spike-in controls (Fig S3A), it demonstrates our baits are specific and sensitive to any changes in the quantity of targeted sequences. To test the reproducibility of each biological replicate, we generated a sample correlation plot (Fig 2A). Results of three biological replicates from the same combination of genotype and treatment group together based on their similarities, and the average pair-wise correlation between them within groups is above 80% (Fig 2A). Thus, the RNA-CAP-I-seq method is highly repeatable. To check how well our RNA-CAP-I-seq captured differential gene expression, we visualized the mapped reads in a genome browser. The overall expression pattern of *SARD1* gene in three biological replicates under all five different treatments is similar (Fig 2B). More reads were mapped to *SARD1* in the samples from “PTI”, “PTI + t-ETI” and “PTI + c-ETI” than those in mock or untreated samples, which is consistent with the previous observation of *SARD1* as one of the ERGs from the total RNA-seq data [9]. Pathogen-induced SA accumulation is required for plant immunity, and one major pathway of SA biosynthesis is *via* isochorismate (IC) [28]. The IC pathway involves several enzymes that are required for the key catalytic steps, and encoded by *ICS1*, *EDS5* and *PBS3* [29, 30]. They are all ERGs and directly regulated by TFs SARD1 and CBP60g [9, 23]. These three SA biosynthetic genes are usually transcriptionally co-regulated in the activation of immunity and are also all highly induced in our ‘PTI’ and ‘PTI + ETI’ samples (Fig 2C). Furthermore, ‘PTI + ETI’ induces stronger expression of these genes than ‘PTI’ alone (Fig 2C), potentially through the regulation of SARD1 and CBP60g. In contrast, the transcripts of the house-keeping genes, *UBQ10* and *ACT7* are stable regardless of the treatments (Fig 2D).

**Figure 2.**
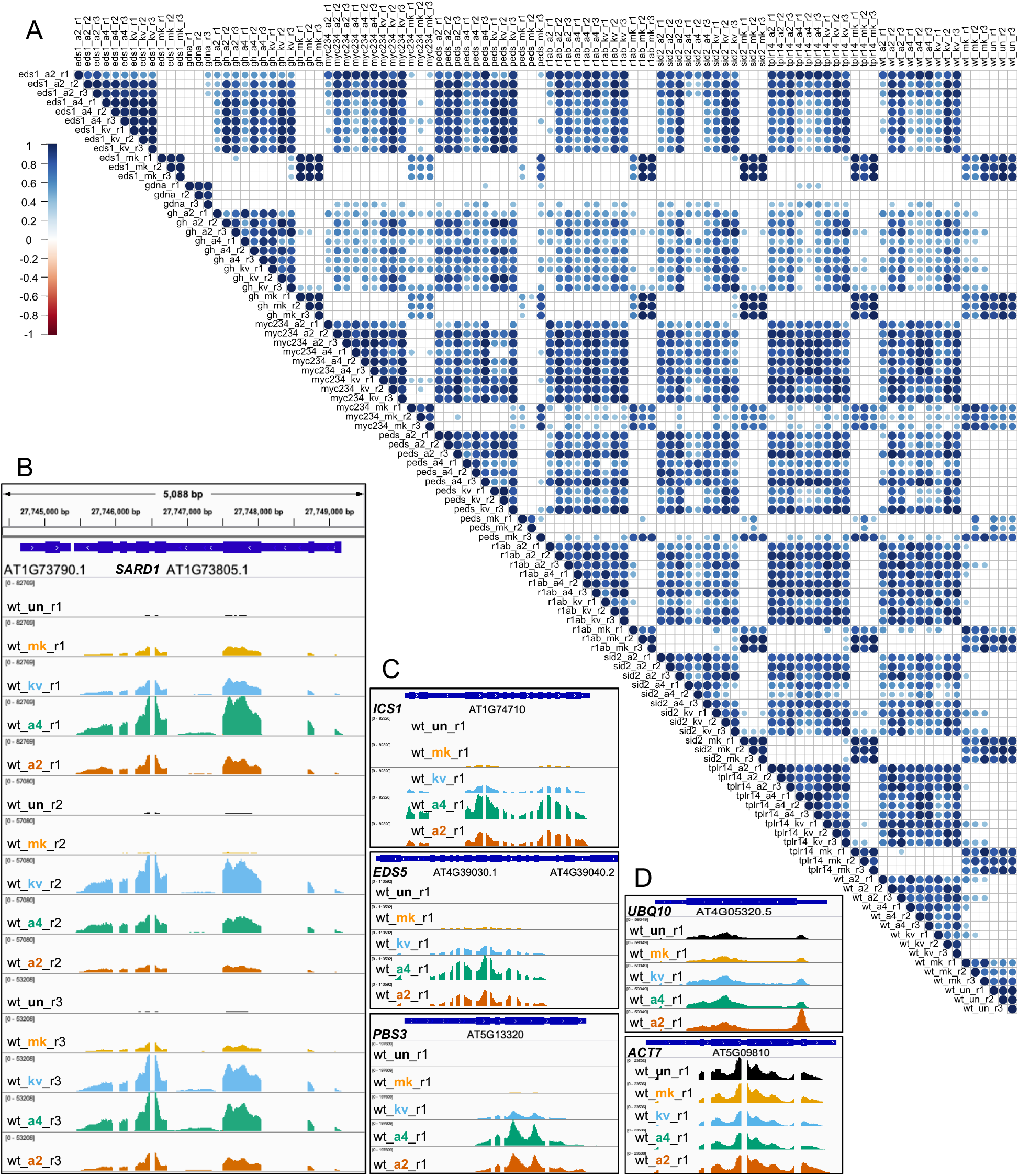
Reproducibility Test of RNA-CAP-I-seq. (A) Correlation analysis of mapped reads from all individual libraries from RNA-CAP-I-seq. All individual libraries including cDNA libraries and spiked-in gDNA libraries from the same CAP-I-seq are pair-wisely compared. 1 indicates 100% positive correlation based on the distribution of reads, while −1 indicates 100% negative correlation. (B) to (D) Mapped reads before normalization are visualized in several CAP-I gene loci in a genome browser. (B) Visualization of reads mapped to *SARD1* locus from wt samples. All three biological replicates (r1-r3) of wt plants under five different treatments are visualized in IGV genome browser at *SARD1* locus. Black indicates untreated (un); orange indicates samples collected at 4 hour post infiltration (hpi) of mock (10 mM MgCl_2_) treatment (mk); sky blue indicates samples collected at 4 hpi of Pf0-1:AvrRps4^KRVYmut^ (kv); bluish green indicates samples collected at 4 hpi of Pf0-1:AvrRps4 (a4); vermilion indicates samples collected at 4 hpi of Pf0-1:AvrRpt2 (a2). See also Figure S2.

Though we observed what we expected from the mapped reads, they required normalization for statistical analysis of relative gene expression. For this, we have developed an R package to normalize and visualize the data generated with sequence capture [31]. From the parameter of ‘Goodness Of Fit’, we found that not all selected control genes are suitable for normalization as some of them are highly variable across 99 samples (Fig S3B). After normalization, we obtained a balanced read distribution with low variation across all samples (Table S4 and S5), enabling statistical analysis for differential gene expression. In the clustering analysis, we retrieved three main clusters of genes based on their expression patterns in all 32 different treatments compared to untreated Col-0 samples (Fig 3A, Table S6). The majority of ERGs are in Cluster I and mostly are immunity related, while Cluster III comprises predominantly control genes (Fig 3B, Table S7). Cluster II contains equal numbers of ERGs and control genes (Fig 3A and 3B). From the same analysis, we also identified three groups of conditions categorizing combinations of genotypes and treatments. Regardless of the genotype, all mock treated samples are clustered in Group I with similar expression patterns of CAP-I genes, indicating they serve as a good negative control for other treatments. In Group III, overall expression of CAP-I genes had no discernable pattern compared to that in Group I and II. In Group II, we were able to identify mutants that have greater impacts on ERG expression pattern in response to treatments (Fig 3A). All Pf0-1-treated samples in *sid2* mutant exhibit similar expression profiles, as do those in *sard1 cbp60g* double mutant. These indicate that ICS1 or SARD1/CBP60g are required for the activation of both ‘PTI’ and ‘PTI + ETI’. Consistent with EDS1 being required for AvrRps4- but not AvrRpt2-induced ETI, our results also show that ERGs in *eds1* are induced less by Pf0-1:AvrRps4 and Pf0-1:AvrRps4^KRVYmut^ (eds1_a4 and eds1_kv) in comparison to those induced by Pf0-1:AvrRpt2 (eds1_a2) (Fig 3A). We also observed that ERGs are induced less in a *pad4 ein2 dde2 sid2* quadruple mutant (*peds*) than in wt by ‘PTI’, which is consistent with previous reports [27, 32]. However, we did not see a strong ERG difference between *peds* and wt in response to ‘PTI + ETI’ (Fig 3A).

**Figure 3.**
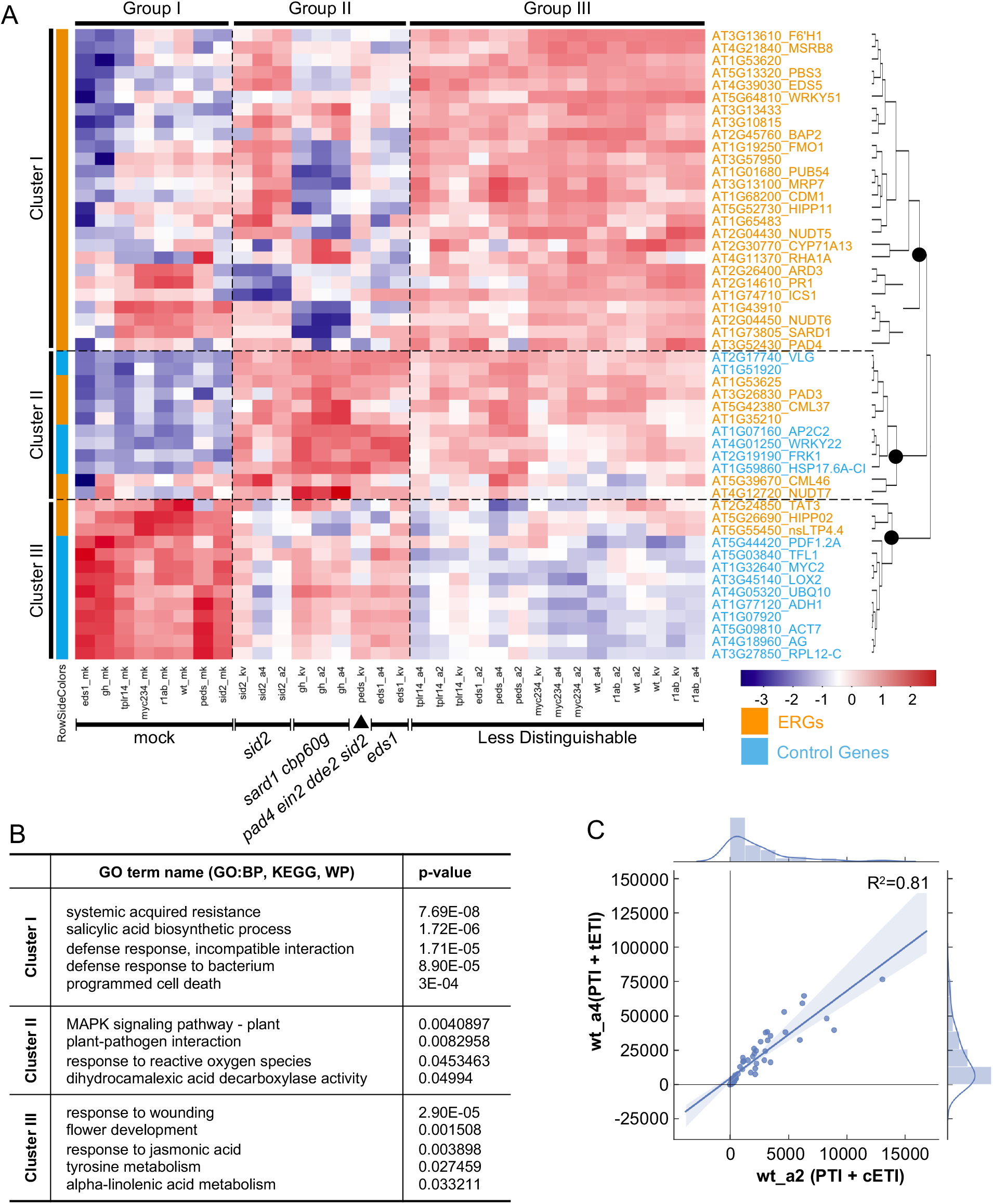
Quantification of RNA-CAP-I-seq. (A) Cluster analysis of normalized read counts from each combination of conditions in comparison to untreated wt Col-0 samples (wt_un). Each combination of conditions represents all combinations of each genotype (wt, eds1, r1ab, sid2, gh, myc234, tplr14, peds) with each treatment (mk, kv, a4, a2). CAP-I genes form three major clusters based on their expression patterns cross all conditions. All conditions form three major groups based on their overall differential gene expression of CAP-I genes. ERGs from CAP-I are in orange and control genes are in sky blue. Heatmap is based on mean z-scores of three biological replicates. Redder color indicates a higher value of z-score, while bluer means a less value of z-score. (B) Top hits of gene ontology (GO) terms based on their p-values for CAP-I genes in each cluster from (A). BP stands for biological process, KEGG is based on the database from Kyoto Encyclopedia of Genes and Genomes. WP refers to WikiPathways database. (C) Comparison of differential gene expression patterns of all CAP-I genes activated by ETI between RRS1/RPS4 and RPS2 in addition to PTI. See also Figure S3, Table S4 to S7.

t-ETI and c-ETI confer resistance *via* different types of NLRs and signaling components [8, 20]. However, there is no previously reported side-by-side comparison of TIR-NLR- and CC-NLR-induced genes upon NLR activation. Here, we compared the induction patterns of ERGs in wt treated with ‘PTI + t-ETI’ and ‘PTI + c-ETI’, and they significantly resemble each other for all CAP-I genes (R^2^=0.81) (Fig 3C). As the 32 conditions are combinations of both genotypes and treatments, we checked the correlation of gene expression patterns with either genotypes or treatments separately (Fig 4A). Gene expression patterns from the treatments of ‘PTI + t-ETI’ and ‘PTI + c-ETI’ within the same genotype tend to group together, rather than with ‘PTI’ (Fig 4A), which further proves that gene expression patterns induced by TIR-NLRs and CC-NLRs at early immune activation stages are similar.

**Figure 4.**
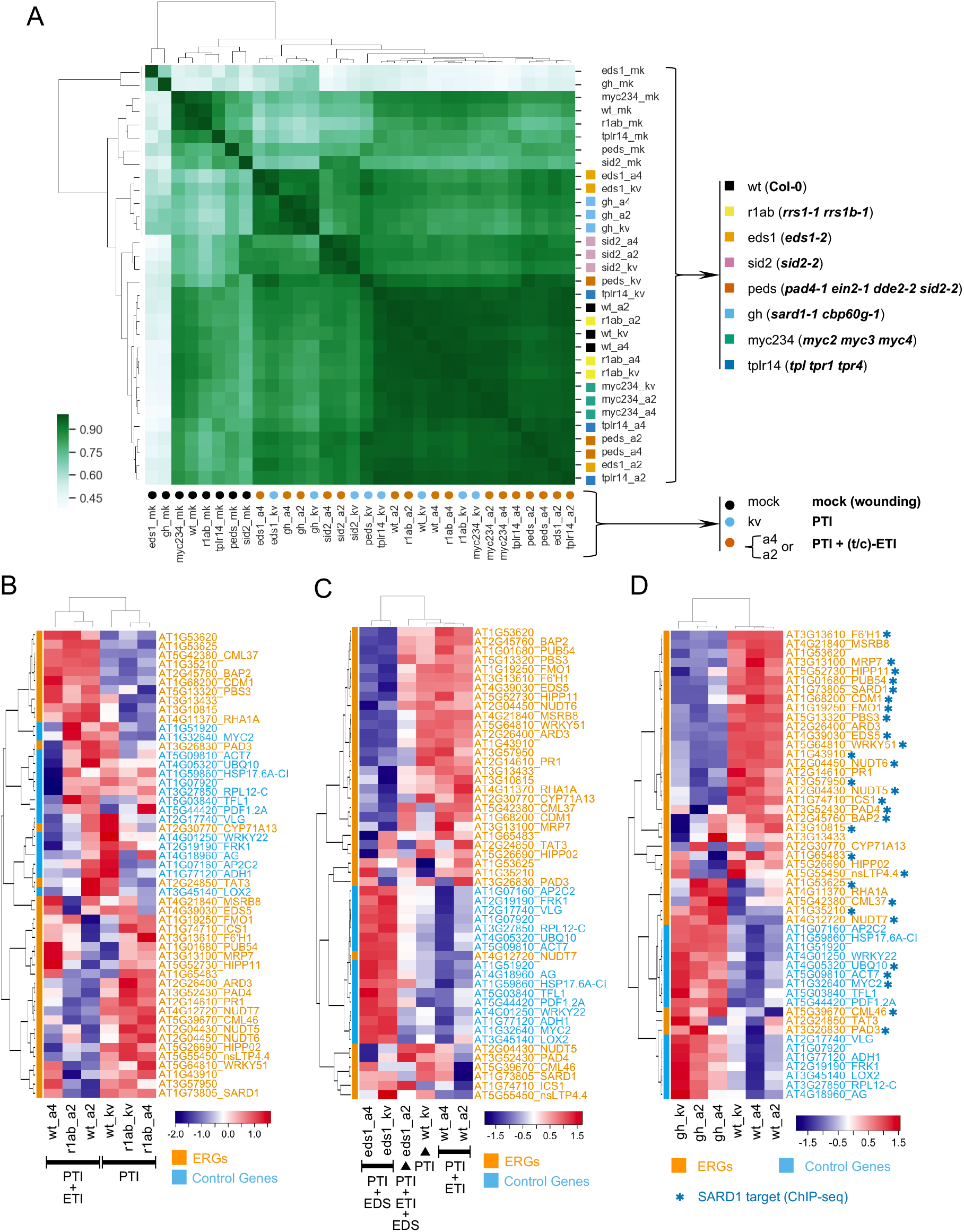
Correlation studies of RNA-CAP-I-seq from different genotypes and treatments. (A) Correlation analysis with mapped and normalized reads from 32 different combinations of both genotypes and treatments. For treatments, we use color-filled circles to indicate, Black circles stand for mock treatment. Sky blue circles are for Pf0-1:AvrRps4^KRVYmut^ (kv). Vermilion circles are for Pf0-1:AvrRps4 or AvrRpt2. For genotypes, we use color-filled squares to indicate, Black squares are for wt Col-0. Yellow squares are for *rrs1-1 rrs1b-1* double mutants. Orange squares are for *eds1-2* (Col-0) mutant. Reddish purple squares are for *sid2-2* mutant, Vermilion squares are for *pad4-1 ein2-1 dde2-2 sid2-2* quadruple mutants. Sky blue stands for *sard1-1*. Bluish green stands for *myc2/3/4, Blue* is for TOPLESS mutants *tprl tpr1 trpr4*. (B) to (D) Differential gene expression are visualized with heat maps. (B) Heatmap of differential expression of CAP-I genes in *rrs1 rrs1b* double mutants compared to wt. (C) Heatmap of CAP-I genes in *eds1* mutant compared to wt. (D) Heatmap of CAP-I genes in sard1 cbp60g mutants compared to wt. See also figure S4.

We examined differential gene expression between each individual mutant and wt. As expected, in both *eds1* and *rrs1 rrs1b* mutants, gene expression patterns are similar between the two treatments of Pf0-1:AvrRps4 and Pf0-1:AvrRps4^KRVYmut^, because both EDS1 and RRS1/RRS1B are required for AvrRps4-induced ETI. Loss-of-function of the AvrRps4 receptors (*rrs1 rrs1b*) or the downstream signaling component EDS1 (*eds1*) resemble the loss-of-recognition of AvrRps4 due to the mutation of AvrRps4 (Pf0-1:AvrRps4^KRVYmut^) in wt plants (Fig 4B and 4C). On the other hand, EDS1 and RRS1/RRS1B are not required for AvrRpt2 recognition, so Pf0-1:AvrRpt2 can still induce both PTI and ETI in *eds1* and *rrs1 rrs1b* mutants (Fig 4B and 4C).

The TFs SARD1 and CBP60g bind to the promoters of defense genes to regulate their expression [13, 23]. We observed that most ERGs that are downregulated in *sard1 cbp60g* mutants are also identified as targets of SARD1 from chromatin immunoprecipitation followed by sequencing (ChIP-seq) of SARD1 (Fig 4D) [23].

The *sid2* mutant is known to have no expression of the *ICS1* gene and compromised SA accumulation induced by pathogens, so we expected to see that SA-induced genes were also downregulated. We observed that genes induced by SA and upregulated during SAR, specifically *PR1 and Acireductone Dioxygenase 3* (*ARD3*) were both downregulated in *sid2* (Fig S4A). *SARD1* is also downregulated in *sid2*, indicating that SARD1-dependent regulation of *ICS1* and SA biosynthesis can in turn positively regulate *SARD1* gene expression. TF WRKY51 and its homolog WRKY50 positively regulate SA signaling and negatively regulate JA signaling [17]. In *wrky50 wrky51* loss-of-function mutants, *Plant Defensin 1.2A* (*PDF1.2A*) is downregulated in response to JA [17]. Here, we found in a *sid2* mutant, *WRKY51* is downregulated, while *PDF1.2A* is upregulated (Fig S4A), which is consistent with the negative expression association between *WRKY51* and *PDF1.2A*. In addition, we found *Cytochrome P450 Monooxygenase 71A13* (*CYP71A13*) is downregulated in *sid2* upon activation of innate immunity, indicating that SA might play positive regulatory roles in camalexin synthesis [33].

The expression of JA response genes *Tyrosine Aminotransferase 3* (*TAT3*) and *Lipoxygenase 2* (*LOX2*) but not *PDF1.2A* is positively regulated by MYC2 and its homologues MYC3 and MYC4 [25, 34]. In our RNA-CAP-I-seq data, we found *MYC2, TAT3* and *LOX2* are downregulated in *myc2 myc3 myc4* triple mutants, whereas *PDF1.2A* is upregulated in the triple mutant in response to activation of innate immunity (Fig S4B).

TOPLESS mutants *tpr1 tpl tpr4* show enhanced susceptibility to bacteria DC3000 and DC3000 carrying AvrRps4 [26]. However, this cannot be simply explained by the expression pattern of ERGs, as we found no clear reduction of ERGs in *tpr1 tpl tpr4* mutants (Fig S5C). Previously TOPLESS proteins were reported as transcriptional co-repressors, but there is only slight evidence in our data of TOPLESS repressor activity towards a few specific genes. Here, we found some defense-related ERGs are downregulated, while others are upregulated, in response to both ‘PTI + t-ETI’ and ‘PTI + c-ETI’ compared to ‘PTI’, which indicates that TOPLESS proteins may play dual functions or indirect roles in regulating ERGs. As there is no ChIP-seq data of TOPLESS proteins or related histone modification marks available, their functions remain unclear. Our data, together with previous reports, nevertheless indicate a complex contribution of TOPLESS proteins in regulating genes induced during plant immunity (Fig S4C) [26].

The *peds* mutant carries mutations in genes from four major immune sectors: PAD4 (*pad4*), ethylene (*ein2*), JA (*dde2*) and SA (*sid2*) [27]. We observed that PAD4, SA and JA response genes are downregulated in *peds*, including *PAD4, ICS1*, *EDS5*, *WRKY51*, *CYP71A13*, *MYC2*, *TAT3* and *LOX2* (Fig S4D). It has been reported that the *PEDS-*represented phytohormone network is required for achieving higher amplitude of transcriptional reprogramming during early CC-NLR-activated ETI in addition to PTI than during PTI alone [35]. However in that report [35], the authors used DC3000 instead of Pf0-1 in our case, which can not only trigger ‘PTI + ETI’ but the background effectors in DC3000 can also trigger effector-triggered susceptibility (‘ETS’), so our results using Pf0-1 are ‘cleaner’. We showed a greater expression difference of ERGs activated by ‘PTI’ and by ‘PTI + ETI’ in *peds* mutant compared to wt (Fig S4D). Like AvrRpt2, AvrRpm1 is also recognized by a CC-NLR, Resistance to *P. syringae* pv *maculicola* 1 (RPM1) and activates ETI [8, 36]. Unlike AvrRpt2-induced ETI, AvrRpm1-induced ETI does not require *PEDS-*represented phytohormone network to achieve a high-amplitude transcriptional reprogram within the early time window of ETI activation [35]. Data from the same report indicate that *RPS2*, but not *RPM1*, gene expression is highly reduced in *peds* when ETI was activated [35]. From this we hypothesize that *RPS2* gene expression might be regulated through these four sectors, explaining why all AvrRpt2-induced ERGs are delayed in contrast to AvrRpm1-induced ETI.

Here, using a limited subset of genes (CAP-I), we could distinguish gene expression profiles during ‘PTI’, ‘PTI + c-ETI’, ‘PTI + t-ETI’ in various mutants, particularly the immune gene regulatory components EDS1, ICS1 and SARD1/CBP60g. Inclusion of additional innate immunity genes in the bait library should enable us to distinguish mutants with enhanced resolution. In addition, as all steps for CAP-I are easy to follow and reproducible, CAP-seq can be further implemented in an automated platform for more high-throughput applications.

Single cell RNA-seq (scRNA-seq) for signature genes is available for some plant tissues [37, 38], and could be combined with capture-seq. A set of 100 marker genes has been defined for *Arabidopsis* that can be used to predict the total transcriptome for each species [39]; these could be incorporated into future capture-seq bait library design. Capture-seq is also capable of comparing the changes in the abundance of any DNA sequences, so it is not limited to cDNA libraries, but can be used in other types of DNA libraries, such as ChIP-seq and Assay for Transposase-Accessible Chromatin using sequencing (ATAC-seq) [40, 41]. Finally, capture-seq could also be used to investigate expression of specific pathogen genes during host colonization (Pathogen Enrichment Sequencing: PenSeq) [6, 7]. In summary, sequence capture provides an extremely versatile and cost-effective method to investigate changes in expression of any designated gene set.

## Supplemental Information

**Figure S1.**
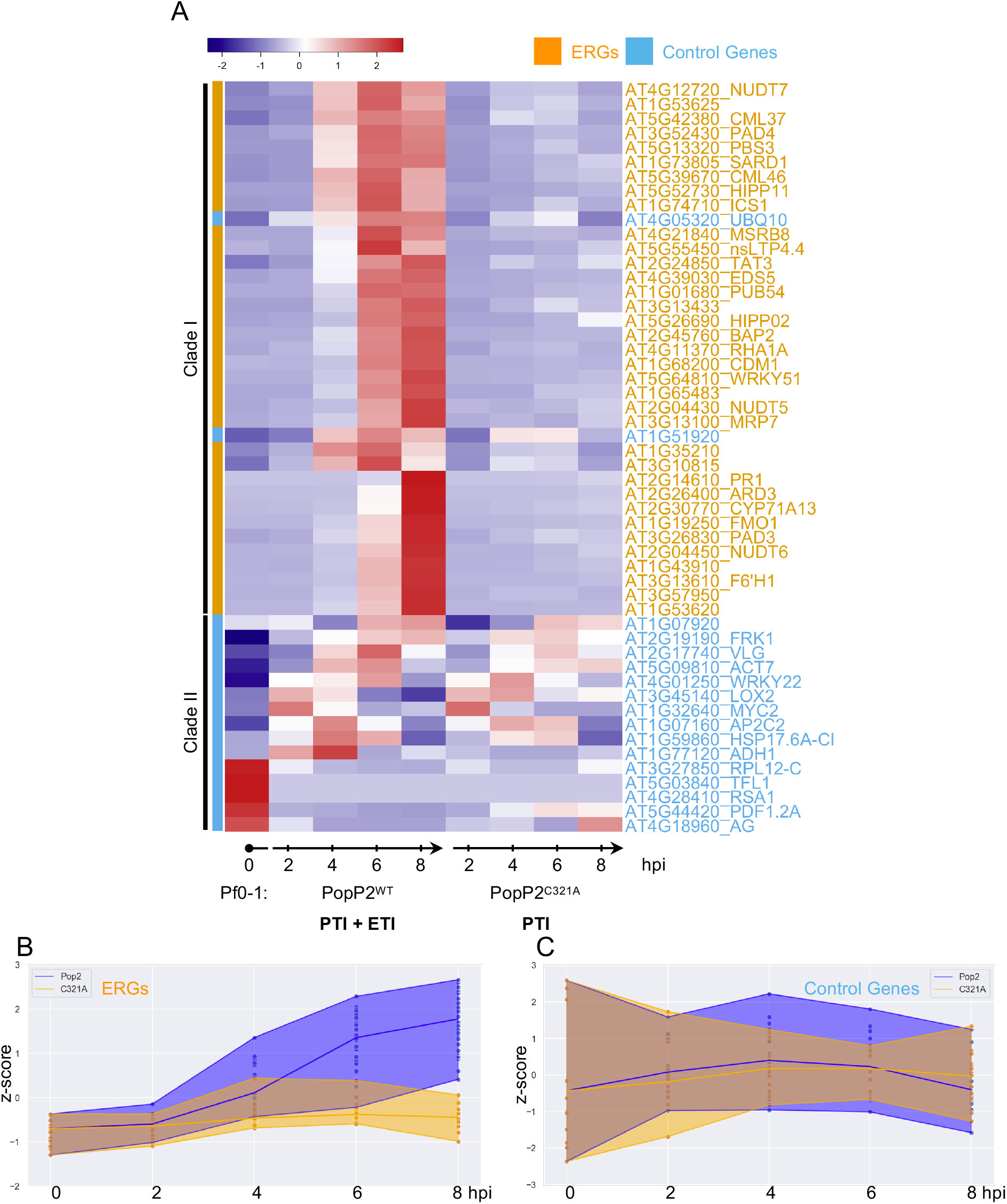
Time-series expression of CAP-I genes under different conditions of immune activation. (A) Heatmap of z-scores for CAP-I genes based on their time-series read counts in the total RNA-seq. ERGs are in orange and selected control genes are in sky blue. Based on the clustering analysis, all ERGs are grouped in one clade, while control genes are grouped in another clade except for two genes. (B) and (C) Time-series visualization of the mean value of z-scores for ERGs and control genes. (B) ERGs show overall upregulations under PTI + ETI treatment compared to PTI alone. (C) Control genes didn’t show such pattern as (B), but overall are stable in their expression level during the course of both immune activations.

**Figure S2.**
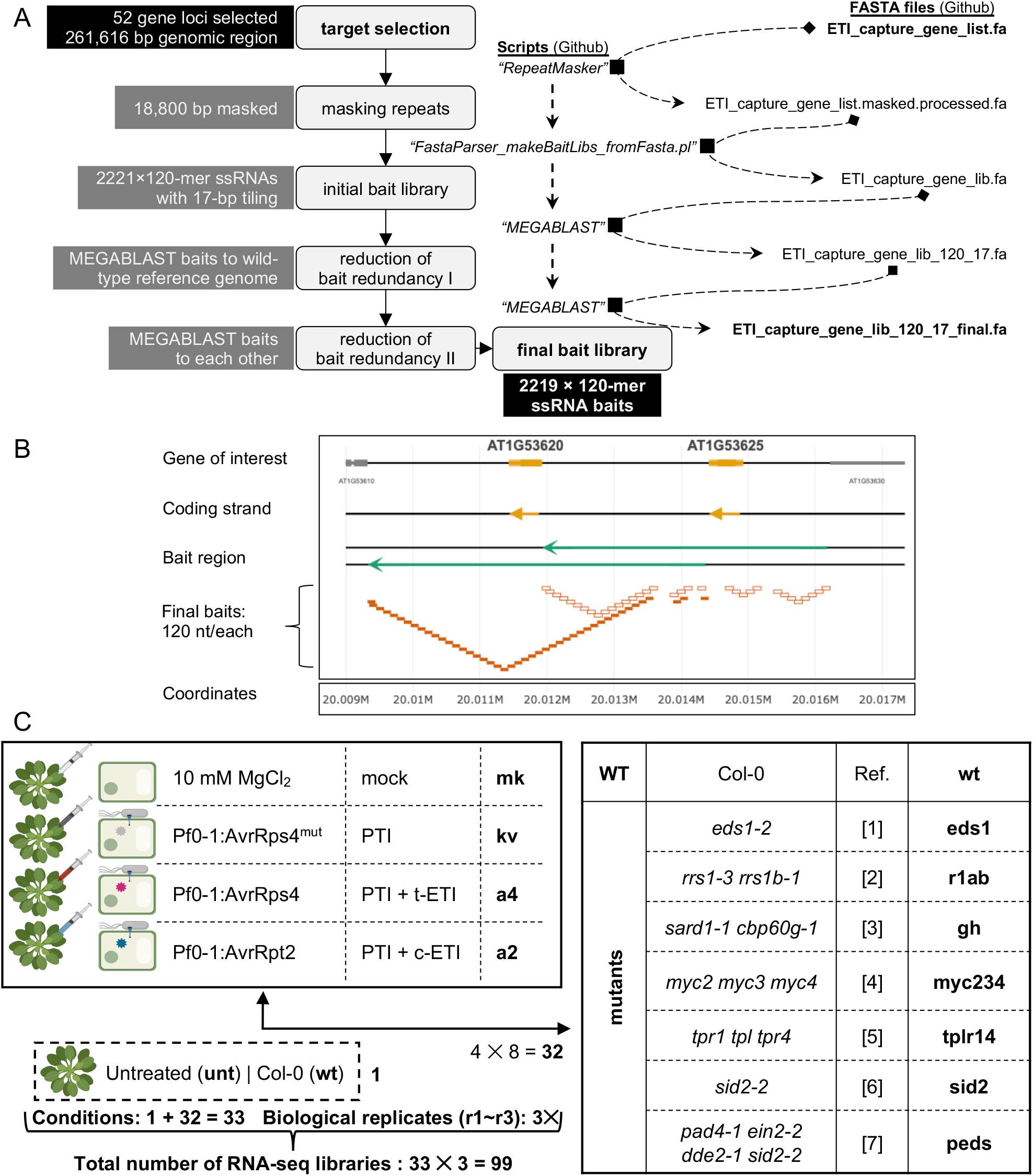
CAP-I bait design and RNA-CAP-I-seq experimental design. (A) Pipeline for CAP-I bait library design. All script information and original files are available in our Github via: https://github.com/slt666666/Ding_etal_2019_CAP_I (B) Visualization of one duplicated region for CAP-I bait design. AT1G53620 and AT1G53625 are two ERG loci next to each other in the genome. The orange overlapped squares with filled orange color are baits for AT1G53620, and those squares without filling color are baits for AT1G53625. Because they are neighboring genes on the genome, so we have got two sets of baits for the overlapped region between these two genes. (C) Experimental design for RNA-CAP-I-seq. Plants from eight different genotypes including wild-type Col-0 accession are treated with four different conditions, which generates 32 different combinations. Untreated Col-0 wt plants are included as an additional control condition, so there are 33 different conditions. All combinations have 3 biological replicates for later-on statistics, so in total, we have 99 individual libraries for CAP-I-seq.

**Figure S3.**
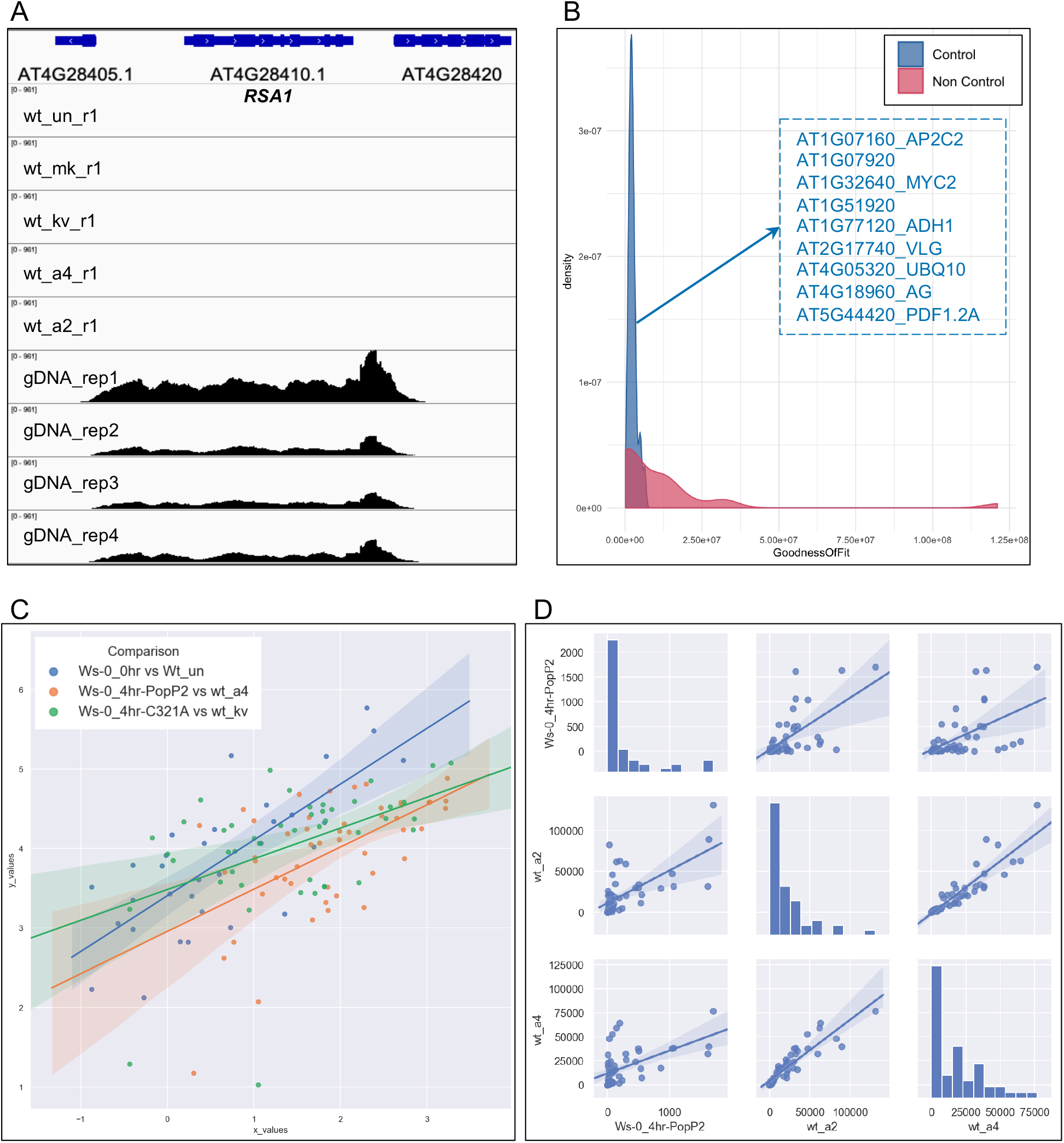
Overall quality assessment of RNA-CAP-I-seq data. (A) RNA-CAP-I-seq reads mapped to control gene *RSA1*. There are nearly no reads mapped to *RSA1* in all 99 cDNA sequencing results, while there are 100% coverage of reads at *RSA1* locus from gDNA. Here we show mapped reads from five different cDNA libraries as examples with the gDNA mapped reads as control. (B) List of control genes for normalizing overlapped with CAP-I control gene set. Density plot with Goodness Of Fit (GOF) shows 9 selected control genes that are much less variable. (C) Comparison of RNA-CAP-I-seq data with previous published RNA-seq data with similar conditions. Ws-0_0hr in previous publication is treated equivalently as untreated wt (wt_un) in this study. Ws-0_4hr-PopP2 stands for 4 hpi of Pf0-1:PopP2 in wt Ws-0, which activates ETI via RRS1-R/RPS4 in addition to PTI; this is treated equivalently as wt_a4, as this activates ETI via RRS1-S/RPS4 in addition to PTI. Similarly, C321A stands for mutant PopP2 and kv stands for mutant AvrRps4, in which case, both only activate PTI. The comparison is based on the normalized read counts from both datasets. (D) Pair-wise comparisons of CAP-I gene expressions from AvrRpt4-, AvrRpt2- and PopP2- induced wt samples.

**Figure S4.**
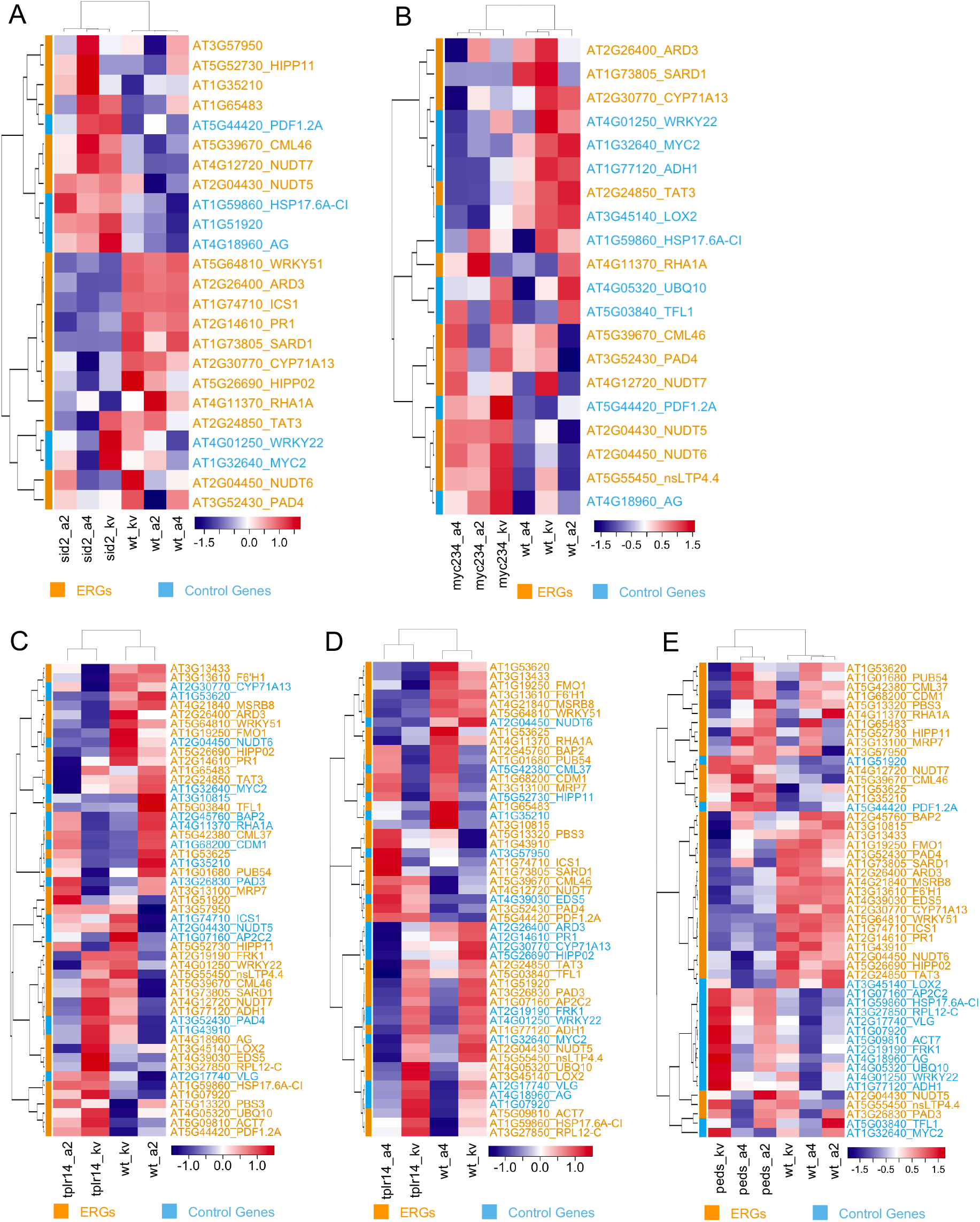
Heatmaps of differential gene expression of mutants individually compared to wt. (A) to (E) are heatmaps based on z-scores, they are for *sid2*, *myc2/myc3/myc4*, *tpr1 tpl tpr4* and *pad4 ein2 dde2 sid2* individually compared with wt.

**Table S1 Information_of_ERGs_and_Control_Genes_in_CAP-I**

**Table S2 Time-series_Expression_of_CAP-I_genes**

**Table S3 Coverage_Information_of_CAP-I_gDNA_seq_Trial**

**Table S4 Read_Counts_of_RNA-CAP-I-seq_before_Normalisation**

**Table S5 Read_Counts_of_RNA-CAP-I-seq_post_Normalisation**

**Table S6 Log_Matrix_of_RNA-CAP-I-seq_Normalised_to_wt_un**

**Table S7 Geno_Ontology_Information_for_Clusters_in_Differential_Gene_Expression_Heatmap**

**Table S8 Barcode_Information_for_RNA-Cap-I_Seq**

### Plant material and growth condition

Mutants of *rrs1-3 rrs1b-1*, *eds1-2*, *sid2-2*, *sard1-1 cbp60g-1*, *myc2 myc3 myc4*, *tpr1 tpl tpr4* and *pad4-1 ein2-1 dde2-2 sid2-2* that were used in this study have been previously described [13, 19, 25–27, 42, 43]. Seeds were sown on compost and plants were grown at 21°C with 10 hours under light and 14 hours in dark, and at 70% humidity.

## METHOD DETAILS

### Bacterial infiltration assay and sample collection

All Pf0-1 strains with different effectors were streaked from their glycerol stock in −70°C freezer on petri dish plates with King’s B medium containing antibiotics for positive selection. Pf0-1:AvrRps4 and Pf0-1:AvrRps4^KRVYmut^ positive colonies were selected with 5 μg/ml tetracycline, 10 μg/ml chloramphenicol and 20 μg/ml gentamycin. Pf0-1:AvrRpt2 were selected with 5 μg/ml tetracycline, 10 μg/ml chloramphenicol and 10 μg/ml kanamycin. Plates were growing in 28°C thermo incubator overnight. Fresh bacteria were streaked off from the plate surface with 1ml clean pipette tips and resuspended in freshly prepared sterile 10 mM MgCl_2_, and spun with 5, 000 rpm for 3 minutes at room temperature. Discarded the supernatant and resuspended the pellet with 10 mM MgCl_2_. The concentration of bacteria was measured and indicated with the optical density at a wavelength of 600 nm (OD_600_). Final concentration of OD_600_=0.2 were used for infiltration with 1 ml needleless syringes. 2 fully expanded leaves from a 5-week-old plant were infiltrated with one of the bacterial strains or just 10 mM MgCl_2_ resuspending buffer as mock. Six leaves from three plants were collected at 4 hours post infiltration (hpi) for each genotype under one certain treatment. Leaves are snap frozen in liquid nitrogen for following up RNA extraction. Three batches of plants were grown under the same condition but on different dates, and samples collected from these three batches are used as three biological replicates.

### RNA extraction

All samples were kept in −70°C freezer from liquid nitrogen if the RNAs were not extracted immediately after sample collection. Total RNAs were extracted with Quick-RNA™ Plant Miniprep Kit (Catalog No. R2024, Zymo Research) following the protocol provided by Zymo Research. The quantities of RNAs were measured by Nanodrop and the qualities of RNAs were assessed with the RNA 6000 Nano Kit (Catalog No. 5067-1511) on an Agilent 2100 Bioanalyzer System. mRNAs were purified with 2 times of enrichment using Dynabeads™ Oligo (dT)25 (Catalog No. 61002, Invitrogen^TM^) from the total RNAs. The qualities and quantities of mRNAs were assessed with the RNA 6000 Pico Kit (Catalog No. 5067-1513, Agilent) on an Agilent 2100 Bioanalyzer System.

### cDNA library construction for RNA-CAP-I-seq

mRNAs were submitted for first strand synthesis with Random Decamers (50 µM) (Catalog No. AM5722G, Invitrogen^TM^) and SuperScript™ IV Reverse Transcriptase kit (Catalog No. 18090200, Invitrogen^TM^). The second strand cDNA synthesis was carried out as previously described [44, 45]. Concentration of double strand cDNAs were quantified with the HS dsDNA Assay kit (Catalog No. Q32851, Invitrogen^TM^) on a Qubit Fluorometer. Illumina sequencing-compatible cDNA libraries were constructed using tagmentation [46]. All libraries were barcoded with in-house custom designed primers (Table S8) and assessed with the High Sensitivity DNA Kit (Catalog No. 5067-4626, Aligent) on an Agilent 2100 Bioanalyzer System.

### CAP-I bait design and RNA-CAP-I sequence capture

For enrichment of selected ERGs and controls, 2219 synthetic 120-nt biotinylated RNA probes with 17 bp tiling were designed and synthesized, complementary to 52 gene regions (including promoter, coding, intron and terminators) totaling 261,616 bp from the reference genome of *Arabidopsis thaliana* Col-0 [47] (MYbaits; MYcroarray now is Arbor Biosciences, MI, USA; https://arborbiosci.com/). Repetitive regions of total 18800 bp within the targeted sequences were masked using RepeatMasker (Smit AFA, Hubley R & Green P. RepeatMasker Open-4.0. 2013-2015), and two highly represented baits with >10 MEGABLAST hits to the TAIR10 reference genome were removed [48]. All detailed information can also be found in our GitHub (Link). In preparation for sequencing, barcoded libraries were sized on the Agilent 2100 Bioanalyzer and then quantified using the Qubit Fluorometer and real-time quantitative PCR (Catalog no. KK4824, Kapa Biosystems). Individual samples were pooled equimolarly. After multiplexing, the RNA-CAP-I library was carried out for sequence capture with CAP-I baits following the protocol provided with blockers specifically for indices with 9 nucleotides. (https://arborbiosci.com/wp-content/uploads/2017/10/MYbaits-manual-v3.pdf)

### RNA-CAP-I-seq on a NextSeq 500 sequencer

The multiplexed libraries were used as input following the NextSeq 500 instrument sample preparation protocol (Catalog no. 15048776, Illumina). With a recommended 1.8‐pM library concentration resulted in clustering density in our instrument (276,000 clusters/mm2). Samples were sequenced on a single flow cell of the NextSeq 500/550 High Output kit (75 cycles), using a 74-cycle (single‐end) configuration. The sequencing run in the NextSeq 500 produced over 600 million single‐end reads with a Q30 ≥ 92.5%.

### Demultiplexing raw data from the NextSeq 500

Raw sequence data obtained from Illumina NextSeq500 sequencing platform are per-cycle base call (BCL) format. As many analysis application tools require per-read FASTQ format files as an input, we need to transform bcl file to fastq. A conversion software by Illumina called bcl2fastq version 2.20.0 (http://emea.support.illumina.com/downloads/bcl2fastq-conversion-software-v2-20.html), was used to demultiplex samples and convert the BCL format to FASTQ format. A sample sheet was prepared following the user guide (https://support.illumina.com/content/dam/illumina-support/documents/documentation/software_documentation/bcl2fastq/bcl2fastq2-v2-20-software-guide-15051736-03.pdf). The sample sheet contains sample identifier and a barcode or a barcode pair (nucleotide bases) and is provided to bcl2fastq for correct demultiplexing of the sample sequence reads. More detail about the command line usage of bcl2fastq tool can be obtained in the user guide. All raw reads post demultiplexing will be open access through the European Nucleotide Archive (ENA) under the accession number of PRJEB34520.

### Mapping reads to genome data, transcript annotation, and profiling of gene expression

The single-end reads for cDNA libraries were mapped to the *Arabidopsis thaliana* Col-0 reference genome (TAIR10) using TopHat v.2.1.1 [49]. Reads from the spike-in genomic DNA were aligned to the reference genome using Bowtie2 v2.2.9[50]. The resulting BAM files were sorted with SAMtools before downstream analysis [51]. With sorted BAM files, all downstream analysis following the pipeline of ‘atacR’ [31]. All the data that we were not able to include in the supplemental materials are available in Github (https://github.com/slt666666/Ding_etal_2019_CAP_I). All scripts and files we generated for this study are available in our Github (https://github.com/slt666666/Ding_etal_2019_CAP_I).

## Acknowledgements

We thank Dr. Zane Duxbury, Dr. Hee-Kyung Ahn and Mr. Samuel Holden for careful reading and suggestions on our manuscript.

We thank the Gatsby Foundation (United Kingdom) for funding to the JDGJ laboratory. PD acknowledges a Future Leader Fellowship supported by the UK Biotechnology and Biological Sciences Research Council (BBSRC), grant number BB/R012172/1; and a Marie Skłodowska-Curie Fellowship supported by the European Union’s Horizon 2020 Research and Innovation Program, grant number 656243. BN was supported by the Norwich Research Park (NRP) Biosciences Doctoral Training Partnership (DTP) from BBSRC, grant number BB/M011216/1. OJF, TS, RKS, DM and JDGJ were supported by the Gatsby Foundation funding to the Sainsbury Laboratory.

## Author Contributions

PD and JDGJ conceptualized the study. Experiments and methods of RNA-CAP-I-seq are designed by PD, and are carried out by PD and BN. CAP-I baits and barcodes are designed by PD and OJF. Data analysis are performed by PD, DM, RKS and TS. The original draft was written by PD, and reviewed and edited by JDGJ, BN, OJF, TS, RKS and DM.

